# Unexpected antidepressant-like effects of temozolomide in a mixed sex-cohort of adult rats: role of hippocampal FADD protein

**DOI:** 10.64898/2026.04.23.720315

**Authors:** Laura Gálvez-Melero, M. Julia García-Fuster

## Abstract

Temozolomide is the gold standard chemotherapeutic agent used in the treatment of glioblastoma multiforme. Yet its pharmacological use has been linked to the emergence of depressive- and/or anxiety-like behaviors, probably through the inhibition of hippocampal neurogenesis. Since prior studies reporting these negative effects were based on prolonged treatment paradigms (i.e., from 2 weeks to up to 6 months), and given the few reports that have included female rodents in their studies, our approach aimed at further characterizing the behavioral effects induced by temozolomide (25 mg/kg, 1 or 2 cycles, 5 days/cycle) in a mixed-sex cohort of adult rats. To do so, rats were scored across time through specific behavioral tests that capture diverse manifestations of affective-like responses (forced-swim, open field, novelty-suppressed feeding and sucrose preference) or cognitive performance (Barnes maze). At the neurochemical level, we ascertained the effects of 2 cycles of temozolomide on hippocampal neurogenesis (neural progenitors with NeuroD) and other potential neuroplasticity targets (i.e., FADD, BDNF). The main results showed that temozolomide induced unexpected antidepressant-like responses in a treatment-duration manner while decreased hippocampal FADD, a neuroplastic marker previously associated with the acute and repeated actions of most antidepressants. These results break the prior dogma linking increased hippocampal neurogenesis with antidepressant-like efficacy, and suggest that other mechanisms of action, such as the one described through the neuroplastic molecule FADD, might be responsible for the antidepressant-like actions of temozolomide, even in the presence of impaired neurogenesis. Our results, in conjunction with the prior data, suggested cycle- and/or length-dependent treatment effects in terms of temozolomide’s antidepressant- vs. depressant-like profile, while proposing a novel biomarker of its treatment response.

## Introduction

Temozolomide is the gold standard chemotherapeutic agent used in the treatment of glioblastoma multiforme, the most common primary malignant brain tumor (recently reviewed by Królikowska et al., 2025). Glioblastoma patients experience high rates of depression (reviewed by Mugge et al., 2020), which might be related to the bad prognostic of the disease and its low life expectancy. However, temozolomide’s mechanism of action might also be involved in the induction of such negative affect, since it does not only impair cell proliferation by inducing cell cycle arrest in the G2/M phase of tumor cells (reviewed by Tomar et al., 2021), but it also affects healthy brain cells, reducing adult hippocampal neurogenesis (Garthe et al., 2009; Nokia et al., 2012; Egeland et al., 2017; García-Cabrerizo et al., 2020). Consequently, a reduction of adult hippocampal neurogenesis might be linked to the development of depressive- and/or anxiety-like behaviors (Duman et al., 1999; Revest et al., 2009; Kheirbek et al., 2012), thus suggesting that glioblastoma treatment with temozolomide might be responsible for inducing negative affect. Although this hypothesis has been extensively supported by many preclinical studies (e.g., Egeland et al., 2017; Pereira-Caixeta et al., 2018; Gan et al., 2019; Dey et al., 2020; Cabrera-Muñoz et al., 2023), other reports have suggested that the reduction of hippocampal neurogenesis was not enough to drive a depressive-like phenotype (Volmayr et al., 2003; Samuels and Hen, 2011; see further discussion in García-Cabrerizo et al., 2020). Within this framework, the main objective of the present study was to further characterize the behavioral effects induced by temozolomide (1 or 2 week cycles of 5 days per week) in a mixed-sex cohort of adult rats, given the few reports that have included female rodents in their studies, and since prior studies linking temozolomide with depressive- and/or anxiety-like responses were based on longer treatment paradigms (i.e., from 2 weeks to up to 6 months).

To do so, the effects of temozolomide were scored in a mixed-sex cohort of adult rats after 1 or 2 cycles of treatment through specific behavioral tests that capture different manifestations of affective-like responses (i.e., forced-swim, open field, novelty-suppressed feeding and sucrose preference; Bis-Humbert et al., 2020; Ledesma-Corvi et al., 2022; Jornet-Plaza et al., 2025). Moreover, the long-term effects of temozolomide on cognitive performance were also evaluated in the Barnes maze (short- and long-term memory) since some reports suggested that the depletion of new neurons might also affect hippocampus-dependent learning memory (e.g., Pereira-Caixeta et al., 2018). Finally, since the hippocampus is a brain region crucial for mood regulation and memory, another objective of the present study evaluated hippocampal neurogenesis (i.e., neural progenitors with NeuroD) and other potential neuroplasticity targets that could be mediating temozolomide’s behavioral responses.

On one hand, we selected BNDF since the activation of this neurotrophic factor via subsequent signaling through TrkB receptors has been described as a key transducer of antidepressant-like effects (e.g., Björkholm and Monteggia, 2016; Casarotto et al., 2021; Madjid et al., 2023) and/or when dysregulated of negative affect (e.g., Mondal and Fatima, 2019; Frycz et al., 2026). On the other hand, FADD is a crucial adaptor of death receptors that can engage apoptosis or survival actions (e.g. neuroplasticity) through the balanced regulation of its total (FADD) and phosphorylated (p-FADD) forms. Activation of monoamine receptors, indirect targets of classic antidepressant drugs, reduced FADD and increased p-FADD and p-FADD/FADD ratio in brain (García-Fuster and García-Sevilla, 2015). Similar FADD regulations were observed for selected antidepressant drugs with different mechanism of action (García-Fuster and García-Sevilla, 2016). These results indicated that inhibition of pro-apoptotic FADD adaptor could function as a common signaling step in the initial activation of monoamine receptors (García-Fuster and García- Sevilla, 2015) and antidepressant-like responses in the brain (García-Fuster and García-Sevilla, 2016), as well as being dysregulated in post-mortem brain samples of major depressive disorder patients (García-Fuster et al., 2014). Interestingly, both BDNF (e.g., Kuipers et al., 2016; Hoirisch-Clapauch, 2022) and FADD (e.g., Hernández-Hernández et al., 2023; Colom-Rocha et al., 2023; Colom-Rocha and García-Fuster, 2025) have been linked with hippocampal neurogenesis regulation, and it is in this context and against this background that we aimed to study them.

## Methods

### Animals

For the present study, a total of 102 Sprague-Dawley adult rats (51 males and 51 females) were used in 3 different experimental procedures as detailed in Fig. 1. All procedures were approved by our Local Bioethical Committee (SSBA 21/2024 AEXP; Conselleria Medi Ambient, Agricultura i Pesca, Direcció General Agricultura i Ramaderia, Govern de les Illes Balears) in accordance with ARRIVE guidelines (Percie du Sert et al., 2020) and following EU Directive 2010/63/EU. All rats were bred in the animal facility at the University of the Balearic Islands, separated at weaning and housed in standard cages (2-4 rats/cage/sex) with unlimited access to food and water under specific environmental conditions (22 °C, 70% of humidity, and a 12:12 h light/dark cycle, lights on at 8:00 AM). Procedures were always performed during the light-period in a mixed-sex cohort of adult rats from litters born at similar times and thus with comparable ages (at least 60 days of age) and body weight ranges (i.e., ∼300 g for males and ∼200 g for females at the beginning of the procedures). All efforts were directed towards minimizing the number of rats used, the number of procedures and their suffering.

**Fig. 1.**
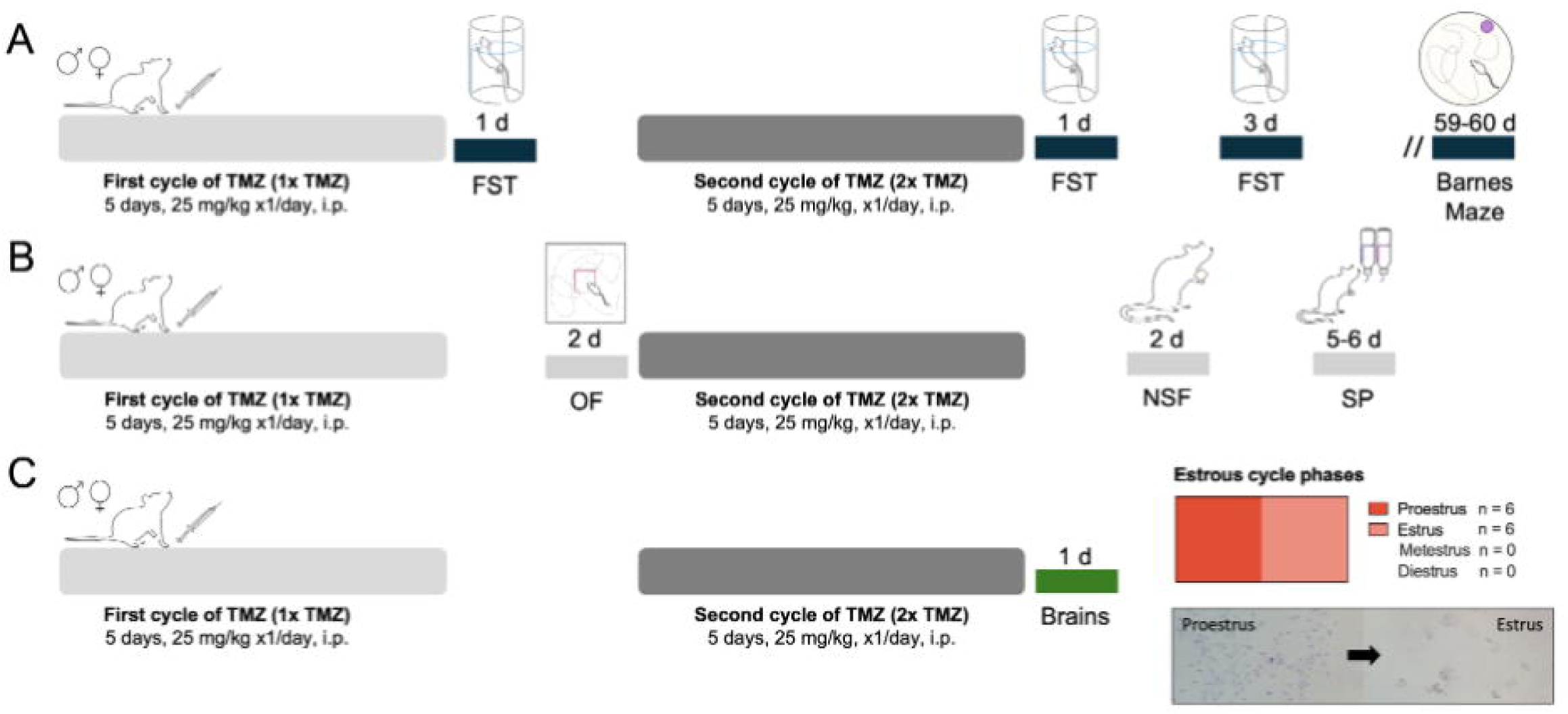
Experimental timeline. **A-B** Behavioral (Studies I and II) and **C** neurochemical (Study III) effects induced temozolomide (TMZ) administered through one- (1x TMZ, 5 days, 25 mg/kg, x1/day, i.p.) or two-cycles (2x TMZ, 10 days total with 2 resting days in between them, 25 mg/kg, x1/day, i.p.) in a mixed-sex cohort of adult rats. A battery of behavioral tests was performed to assess affective- and/or long-term cognitive-like effects produced by TMZ treatment. **A** Study I: forced-swim test, FST. **B** Study II: open field, OF; novelty-suppressed feeding, NSF; sucrose preference, SP; and Barnes maze). **C** Brains were collected to evaluate the effect of TMZ treatment at the neurochemical level as measured 1-day post-treatment (Study III). The estrous cycle phases were evaluated at the time of sacrifice in all female rats.

### Temozolomide drug treatments

Rats were treated with temozolomide (25 mg/kg, i.p.; Merck, Sigma-Aldrich, PHR1437) or vehicle (1 ml/kg, i.p.) for 2 cycles (5 days/cycle, 1 dose/day, 10 days total with 2 resting days in between cycles) as shown in Fig. 1. The selection of temozolomide treatment paradigm (dose and length of treatment) was based on our prior study determining the optimal cyclic temozolomide-treatment conditions needed to reduce basal neurogenic markers in the hippocampus (García-Cabrerizo et al., 2020), as well as in prior studies (e.g., Nokia et al., 2012). While 1 cycle of temozolomide was sufficient to reduce cell proliferation, 2 cycles were needed to reduce early neuronal survival (García-Cabrerizo et al., 2020). This treatment paradigm was followed in 3 separate experiments: Study I (Fig. 1A): Vehicle (Veh, n = 19; 10 males and 9 females); temozolomide (TMZ, n = 20; 10 males and 10 females); Study II (Fig. 1B): Vehicle (Veh, n = 19; 10 males and 9 females); temozolomide (TMZ, n = 21; 10 males and 11 females); Study III (Fig. 1 C): Vehicle (Veh, n = 11; 5 males and 6 females); temozolomide (TMZ, n = 12; 6 males and 6 females). Each experimental design aimed at characterizing the effects of temozolomide behaviorally (Studies I and II) or neurochemically (Study III) as detailed below.

### Forced-swim test

Prior to any drug treatment, basal performance (immobility, climbing and swimming) was scored under the stress of the forced-swim test in adult naïve rats from Study I (Fig. S1) as previously described (e.g., García-Cabrerizo et al., 2020). To do so, all rats were exposed to a 15-min pre-test session in which they were individually placed in water tanks (41 cm high x 32 cm diameter, 25 cm depth; temperature of 25 ± 1 °C), followed, the next day, by a 5-min test session that was videotaped. Videos were blindly evaluated (Behavioral Tracker software, CA, USA) to establish individual basal levels of immobility (i.e., lack of movement except the one required to keep the rat’s nose above the water level) vs. active behaviors (climbing or swimming). These values showed no basal differences between male and female rats and were used to counterbalance experimental groups while ensuring a normal distribution and homogeneity of variance for all groups (see Supplementary Fig. S1).

Remarkably, the forced-swim test is the goal standard screening tool for antidepressants in rodents (Slattery and Cryan, 2012) with predictive validity for its later efficacy in the clinic, which has been slightly adapted for our experimental conditions (see for example, Ledesma-Corvi and García-Fuster, 2022, 2023; Jornet-Plaza et al., 2025; García-Cabrerizo et al., 2025). Thus, the antidepressant-like potential of temozolomide was scored across time by 5-min forced-swim test sessions at the indicated times: 1-day post-1 cycle and 1- and 3-days post-2 cycles (Fig. 1A). These sessions were videotaped to later evaluate the behavioral responses; decreased immobility and increased escaping responses (i.e., climbing and/or swimming) as indicative of an antidepressant-like response with Behavioral Tracker software (CA, USA) under blinded conditions. Unfortunately, two videos of the vehicle group (2 male rats) did not record and/or save in the computer while recording so the n for that group in this test is lower. Also, as previously described in our studies, our set-up based on repeating forced-swim test across days provides reliable measurements of the progress of this behavioral response while avoiding confounding learning effects due to repetition (e.g., García-Cabrerizo et al., 2020, 2025).

### Open field

Rats from Study II were exposed to the open field test 2 days post-1 temozolomide cycle (Fig. 1B). This test measures changes in exploratory-like behavior under the stress of a novel environment. Following prior studies (Bis-Humbert et al., 2020, 2021), the test was done in a wall-enclosed open field arena (85 x 54 cm) under housing illumination conditions (Jornet-Plaza et al., 2025). All rats were initially placed in a corner facing the wall and allowed to freely move and explore the field for 5 min while sessions were recorded. Later, and blinded to the experimental conditions, videos were analyzed with a digital tracking system (Smart Video Tracking software, Version 3, Panlab SL, Barcelona, Spain). Distance travelled (cm), entries to center and time in center (s) were used to ascertain potential anxiety-like changes (Bis-Humbert et al., 2020, 2021).

### Novelty-suppressed feeding

The novelty-suppressed feeding test scores antidepressant-like responses under a stressful situation, generated by food deprivation (48 h) prior to individually placing rats in a square open arena (60 cm x 60 cm, and 40 cm in high) with three food pellets in the center (Bis-Humbert et al., 2020, 2021). Under these conditions, rats from Study II were freely allowed to explore the arena for 5 min under housing lighting conditions (see similar prior procedures: Bis-Humbert et al., 2020, 2021; Ledesma-Corvi et al., 2022; Jornet-Plaza et al., 2025) 2 days after 2-treatment cycles with temozolomide (Fig. 1B). Sessions were videotaped to later blindly analyze latency to food (s), feeding time (s) and food intake (g).

### Sucrose preference test

The sucrose preference test is used to score changes in hedonic-like responses (e.g., Slattery et al., 2007) by allowing rats to select between drinking from a sweet solution (1% sucrose) or water (e.g., Ledesma-Corvi et al., 2022). In our experimental conditions, our goal was to ascertain potential changes induced by temozolomide as measured 5-6 days post-2 treatment cycles. Prior to testing, rats were habituated to drink from two bottles (filled with water) during 24 h placed on each side of the housing cage. Then, for 2 consecutive days (5-6 days post-treatment, Fig. 1B), one of the water bottles was replaced for a bottle containing 1% sucrose, and rats were allowed to freely drink from either one. Bottles were randomly placed in alternate positions in the cage every 24 h to avoid potential preferences to a particular side of the cage.

Results were calculated by weighting the bottles every day to calculate sucrose preference (%), sucrose consumption (ml) and sucrose intake (g/kg) for each rat.

### Barnes maze

The long-term effects of temozolomide (2-treatment cycles) on cognitive performance was evaluated 59- and 60-days post-treatment in rats from Study I in the Barnes maze (Fig. 1B) following standard procedures (i.e., Hernández-Hernández et al., 2018). The Barnes maze we used consists of a circular platform with 18 holes, evenly spaced around its perimeter, and with one of the holes leading to an escape or target box below. Rats conducted the test under visual cues or spatial references to help them locate the escape box, and a bright aversive light stimulus to motivate them to find the target quickly, leveraging their natural agoraphobia. Rats were first habituated to the maze by placing them in a black start chamber located at the center of the maze under a bright light (500 W) for 10 seconds. The chamber was then lifted, and rats were allowed to find and enter the black escape box for up to 3 minutes. Each testing day (59- or 60-days post-treatment; Fig. 1B), rats underwent 3 training trials separated by 10 minutes, which ended either when they entered the target box, or when the time was up. At this point, rats were placed in the target box and left there to habituate for 1 minute. Then, 10 minutes later, the test began, and rats were allowed to find the target box by freely exploring the maze for 90 seconds. The time spent (s) to resolve the maze and the number of errors committed were used as a measure of spatial working memory performance (e.g., Hernández-Hernández et al., 2018; Jornet-Plaza et al., 2025).

#### Sample collection

Brains were collected following rapid decapitation 1 day after the 2 treatment cycles with temozolomide (Study III; Fig. 1C). From each collected brain, the left half-hemisphere was quickly frozen in isopentane (Panreac Química, Barcelona, Spain, cat #143501) at -30 °C, while the right hippocampus was freshly dissected and fast-frozen in liquid nitrogen. All samples collected were stored at -80°C until further us. Moreover, at the time of sacrifice, vaginal fluid was collected for all female rats of this study. This was done with cotton swabs dipped in 0.9% saline and spread it on a slide that was saved at -80 °C until further analysis.

### Estrous monitoring

The specific stages of the estrous cycle were monitored in female rats from Study III at the end of treatment as previously described (García-Cabrerizo et al., 2025). The cyclicity of female rats was not ascertained in Studies I and II to prevent excessive stress only in females while undergoing behavioral phenotyping with their male counterparts. Both sexes seem equally variable due to hormonal periodicity (Becker et al., 2016; Kaluve et al., 2022), in line with our prior study (García-Cabrerizo et al., 2025). In fact, the individual variability observed in this study for each behavioral measure and biological sex reinforced that notion (i.e., homogeneity of variance across sexes as displayed in all figures). The estrous cycle phases were monitored in the slides with vaginal fluid under a light microscope following a cresyl violet staining. Out of the female rats evaluated, the results showed that most of the rats were transitioning between diestrus (12%) to proestrus (76%) or proestrus to estrus (12%) stages, with no rats found in metestrus (0%) (see Fig. 1C). In any case, and as previously reported, both sexes showed similar dispersions across their individual values in all experimental results.

### NeuroD quantification by immunohistochemistry

NeuroD, an intrinsic marker that labels hippocampal early neural progenitors, and known to be decreased by 2 cycles of temozolomide treatment in male rats (García-Cabrerizo et al., 2020), was evaluated in rats from Study III by immunohistochemistry in 30 µm slide-mounted sections covering the whole extent of the left hippocampus (-1.72 to -6.80 from Bregma) as previously described (García-Cabrerizo et al., 2020; Ledesma-Corvi and García-Fuster, 2023; Colom-Rocha and García-Fuster, 2025). To do so, we used 3 slides per animal, each one included every 8th coronal section from the anterior, middle and posterior part of the hippocampus, yielding 8 sections/slide (24 sections/animal). Briefly, tissue was postfixed in 4% paraformaldehyde (Merck, Darmstadt, Germany, cat #76240) and exposed to several steps, such as peroxidase inhibition with 0.1% H_2_O_2_ solution (Thermo Fisher Scientific, cat #426000010) and donkey serum as a blocking agent (Merck, cat #A7906). Then, sections were incubated overnight with goat anti-NeuroD (1:25000; Santa Cruz Biotechnology, CA, USA, cat #sc-1084), followed by a series of sequential incubations; secondary antibody (biotinylated anti-goat, 1:1000, Vector Laboratories, CA, USA, #BA-5000), Avidin/Biotin complex (Vectastain Elite ABC kit; Vector Laboratories, cat #PK-6100), chromogen 3,3’-diaminobenzidine (DAB; Merck, cat #D8001) with nickel chloride (Merck, cat #339350) for signal detection. Sections were then dehydrated in graded alcohols, immersed in xylene (Sharlab, Barcelona, Spain, cat #XI0052) and cover-slipped with Permount® (Thermo Fisher Scientific, cat #SP15-500). Once all slides were dry, they were coded to be blindly quantified. NeuroD +cells were counted in the dentate gyrus region of the hippocampus with a Leica DMR light microscope (63x objective lens with a 10x ocular lens; total magnification of 630x) by focusing across the thickness of the section. The overall number of NeuroD +cells counted for each animal was divided by the hippocampal area quantified (mm^2^) as scanned with a GS-800 Imaging Calibrated Densitomer from BioRad and analyzed with Quantity One Software. This analysis yielded a value of +cells/mm^2^ as previously described (Ledesma-Corvi and García-Fuster, 2023; Colom-Rocha and García-Fuster, 2025).

### Neuroplasticity markers evaluation by western blot

Following standardized procedures (García-Cabrerizo et al., 2020; Ledesma-Corvi and García-Fuster, 2023; Colom-Rocha and García-Fuster, 2025), total homogenates from the right part of the hippocampus were loaded (40 μg) and separated by electrophoresis through 10-14 % SDS-PAGE mini-gels (Bio-Rad Laboratories, CA, USA). Proteins were then transferred to nitrocellulose membranes and incubated (4 °C, overnight) with particular primary antibodies: (1) anti-FADD (H-181) (1:2500; cat #sc-5559; Santa Cruz Biotechnology, CA, USA); (2) anti-BDNF (1:10000; cat #ab108319; Abcam, Cambridge, UK); and (3) anti-ß-actin (clone AC-15) (1:10000; Sigma-Aldrich, MO, USA). The next day, membranes were incubated with specific secondary antibodies linked to horseradish peroxidase (1:5000 dilution; Cell Signaling), ECL chemicals (Amersham, Buckinghamshire, UK), to finally be placed in contact with an autoradiographic film (Amersham ECL Hyperfilm) for 1-60 min. Films were later evaluated by densitometric scanning (GS-800 Imaging Calibrated Densitometer, Bio-Rad). Every mini-gel contained different randomized brain samples from temozolomide- and vehicle-treated groups from both sexes, and each sample was run in at least 3-6 different mini-gels. Percent changes in immunoreactivity were assessed with respect to vehicle-treated samples in each gel (100%), and the mean value was used as a final estimate. Membranes were then stripped and reprobed for ß-actin, which was used as a loading control.

### Data analysis and statistics

Data is shown as a mixed-sex cohort combining both sexes and was analyzed with GraphPad Prism, Version 10 (GraphPad Software, Inc., CA, USA). Results are expressed as mean values ± standard error of the mean (SEM). Individual symbols are shown for each rat (blue symbols represent male rats while orange symbols females). Assumptions for normality of data distribution and homogeneity of variance were met through the Shapiro-Wilk test (i.e., recommended for small sample size <50). All behavioral and neurochemical data was analyzed through unpaired two-tail Student’s *t*-test in a mixed-sex cohort of rats for each treatment group. The level of significance was set at p ≤LJ0.05. Data supporting the present findings will be available upon reasonable request to the corresponding author.

## Results

### Temozolomide induced antidepressant-like responses in a treatment-duration manner

The results demonstrated that the length of temozolomide treatment (1x o 2x cycles, 5 or 10 days in total) was important for inducing changes in the behavioral response observed in the forced-swim test. Particularly, 1x cycle of temozolomide treatment (5 days) did not change the time rats spend immobile (*t* = 0.65, *df* = 35, *p* = 0.519, Fig. 2A), climbing (*t* = 0.62, *df* = 35, *p* = 0.542, Fig. 2B) or swimming (*t* = 0.96, *df* = 35, *p* = 0.346, Fig. 2C).

**Fig. 2.**
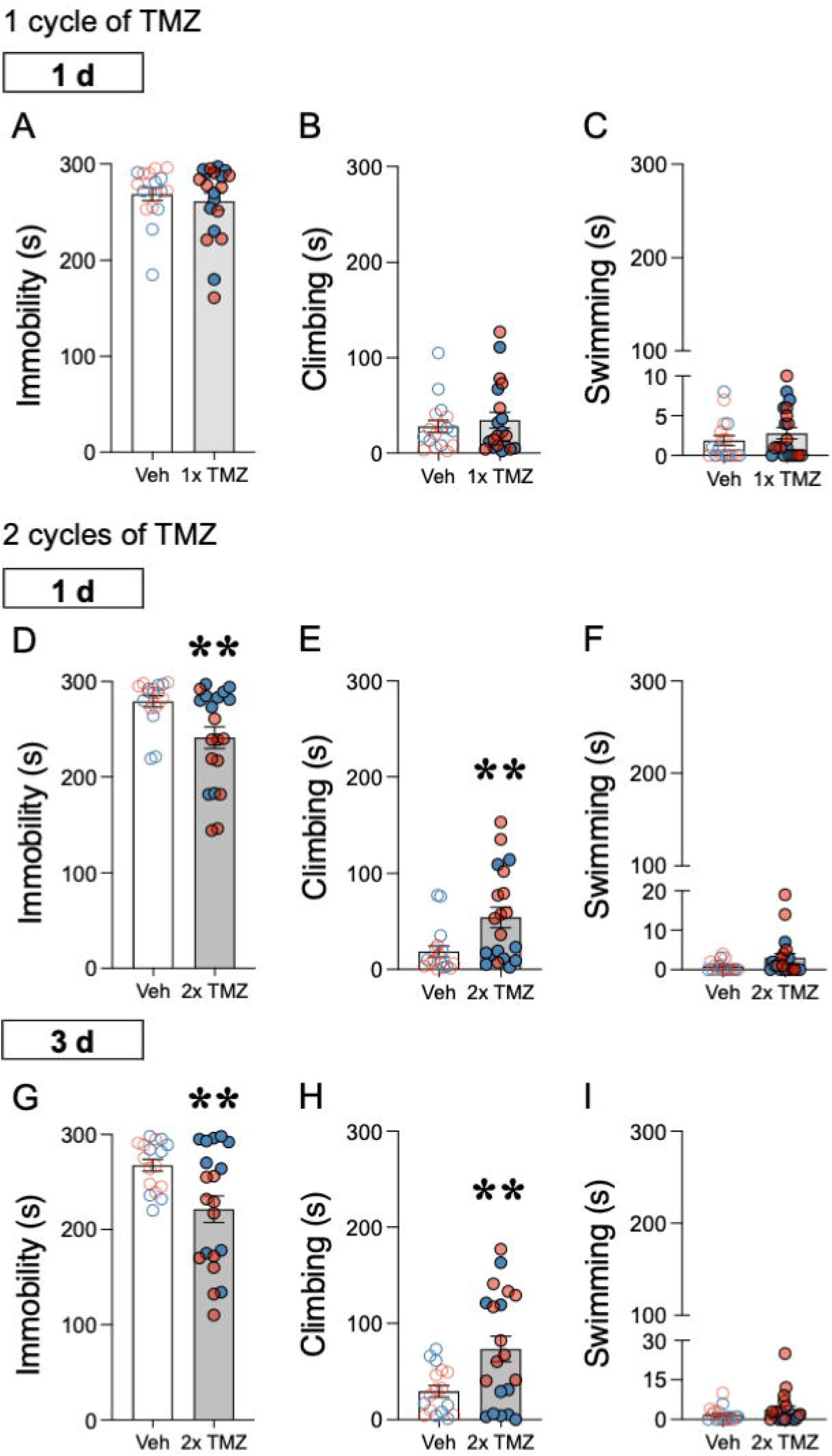
Behavioral effects induced by temozolomide (TMZ) under the stress of the forced-swim test (FST) in a mix-sex cohort of adult rats. **A** Changes in immobility (s), **B** climbing (s), or **C** swimming (s) induced by one TMZ cycle (1x TMZ, 5 days, 25 mg/kg, x1/day, i.p.) or vehicle (Veh) as measured 1-day post-treatment. **D, G** Changes in immobility (s), **E, H** climbing (s), or **F, I** swimming (s), induced by two TMZ cycles (2x TMZ, 10 days total with 2 resting days in between them, 25 mg/kg, x1/day, i.p.) or Veh as measured 1- or 3-days post-treatment. Data represent mean ± SEM of the each of the defined measurements. Individual values are shown for each rat (blue symbols represent male rats while orange symbols females). Data is shown as a mixed-sex cohort combining both sexes. The effect of TMZ treatment was evaluated through Student *t*-tests (\*\**p* < 0.01 when comparing 2x TMZ vs. Veh-treated rats).

Interestingly, and contrary to what was initially expected, 2x cycles of temozolomide treatment (10 days) induced behavioral responses indicative of an antidepressant-like response as observed 1- (Fig. 2D-F) and 3-days (Fig. 2G-I) post-treatment. The results showed an effect of treatment for immobility and climbing at both time points of analysis in a mixed-sex cohort of adult rats. Specifically, 2x cycles of temozolomide induced a significant reduction in immobility as observed 1- (-38 ± 13 s, *t* = 2.81, *df* = 35, \*\**p* = 0.005 vs. vehicle-treated rats; Fig. 2D) and 3-days (-46 ± 16 s, *t* = 2.84, *df* = 35, \*\**p* = 0.008 vs. vehicle-treated rats; Fig. 2G) post-treatment in adult rats. This reduced immobility aligned with increased climbing behavior (1-day: +35 ± 13 s, *t* = 2.78, *df* = 35, \*\**p* = 0.009 vs. vehicle-treated rats; Fig. 2E; 3 days: +46 ± 16 s, *t* = 2.84, *df* = 35, \*\**p* = 0.008 vs. vehicle-treated rats; Fig. 2H), in line with an antidepressant-like response.

### Temozolomide did not modify the behavioral responses observed in a battery of tests used to measure affective-like behavior

The results demonstrated that temozolomide treatment (1x o 2x cycles, 5 or 10 days in total) did not induce changes in other affective-like dimensions other than the ones just described in the forced-swim test in a mixed-sex cohort of adult rats (Fig. 3). This was the case for potential changes in the open field as measured 2-days post-1x temozolomide cycle (Fig. 3A-C), and the novelty-suppressed feeding test (Fig. 3D-F) and the sucrose preference (Fig. 3G-I) as measured 2- and 5-6 days post-2 temozolomide cycles, respectively.

**Fig. 3.**
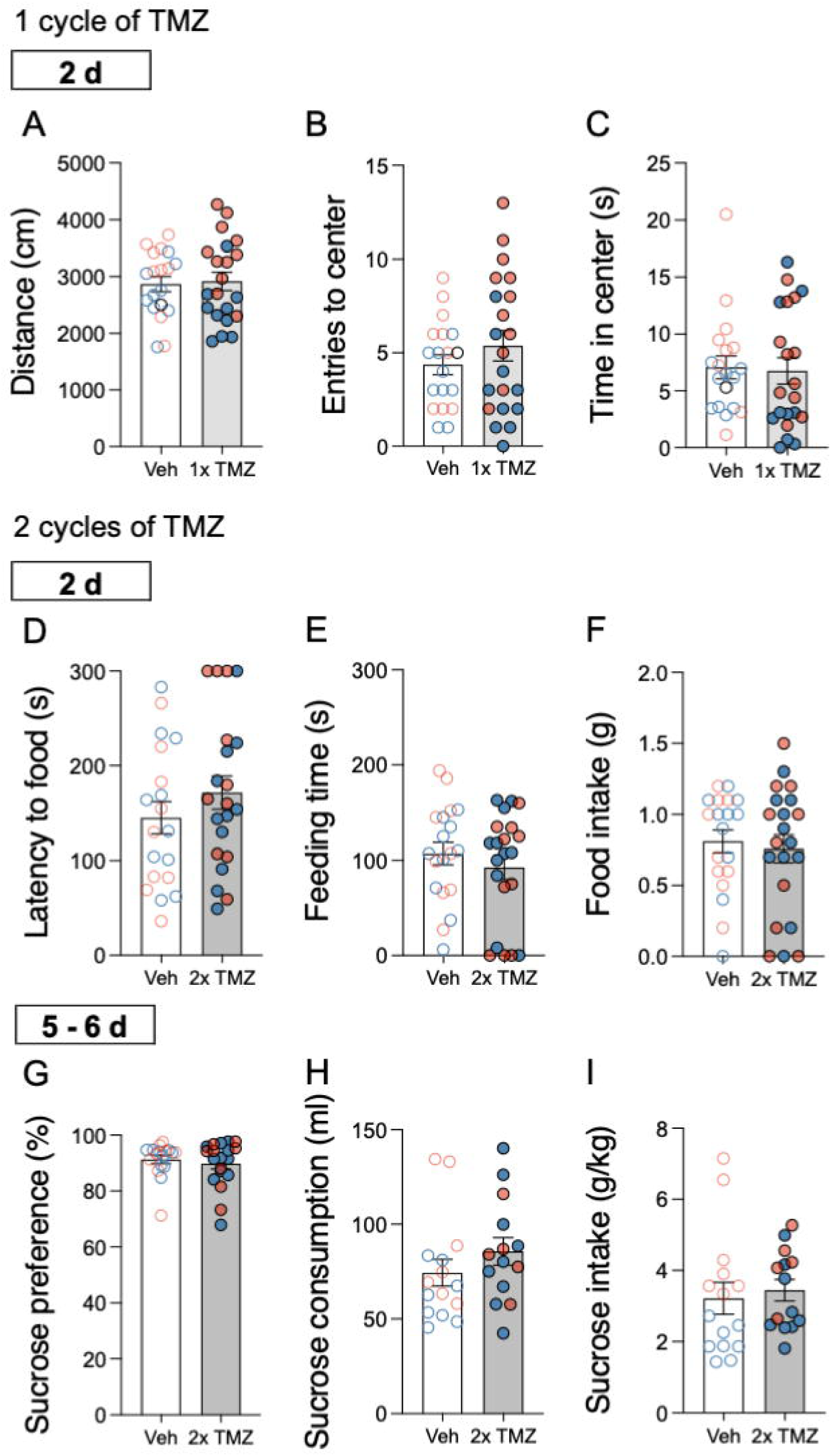
Behavioral effects induced by temozolomide (TMZ) as assessed by a battery of tests across time in a mix-sex cohort of adult rats. **A** Changes in distance (cm), **B** entries to center, or **C** time in center (s) induced by one TMZ cycle (1x TMZ, 5 days, 25 mg/kg, x1/day, i.p.) or vehicle (Veh) as measured 1-day post-treatment in the open field test. **D** Changes in latency to food (s), **E** feeding time (s), or **F** food intake (g) induced by two TMZ cycles (2x TMZ, 10 days total with 2 resting days in between them, 25 mg/kg, x1/day, i.p.) or Veh as measured 2-days post-treatment in the novelty-suppressed feeding test. **G** Changes in sucrose preference (%), **H** sucrose consumption (ml), or **I** sucrose intake (g/kg) induced by two TMZ cycles (2x TMZ, 10 days total with 2 resting days in between them, 25 mg/kg, x1/day, i.p.) or Veh as measured 5-6-days post-treatment in the sucrose preference test (i.e., two-bottle choice). Data represent mean ± SEM of the each of the defined measurements. Individual values are shown for each rat (blue symbols represent male rats while orange symbols females). Data is shown as a mixed-sex cohort combining both sexes. The effect of TMZ treatment was evaluated through Student *t*-tests.

### Temozolomide preserved long-term cognitive function

Cognitive performance was evaluated in a mixed-sex cohort of adult rats from Study I in the Barnes maze 59- and 60-days post-2x cycles of temozolomide treatment (Fig. 4). The results showed that temozolomide preserved the long-term cognitive function of adult rats after a repeated treatment. This was the case when measuring the time (s) and the number of errors committed to resolve the maze both 59-days post-treatment (Fig. 4A-B), and then 24-h later (60-days post-treatment; Fig. 4C-D), demonstrating a preserved spatial working memory performance.

**Fig. 4.**
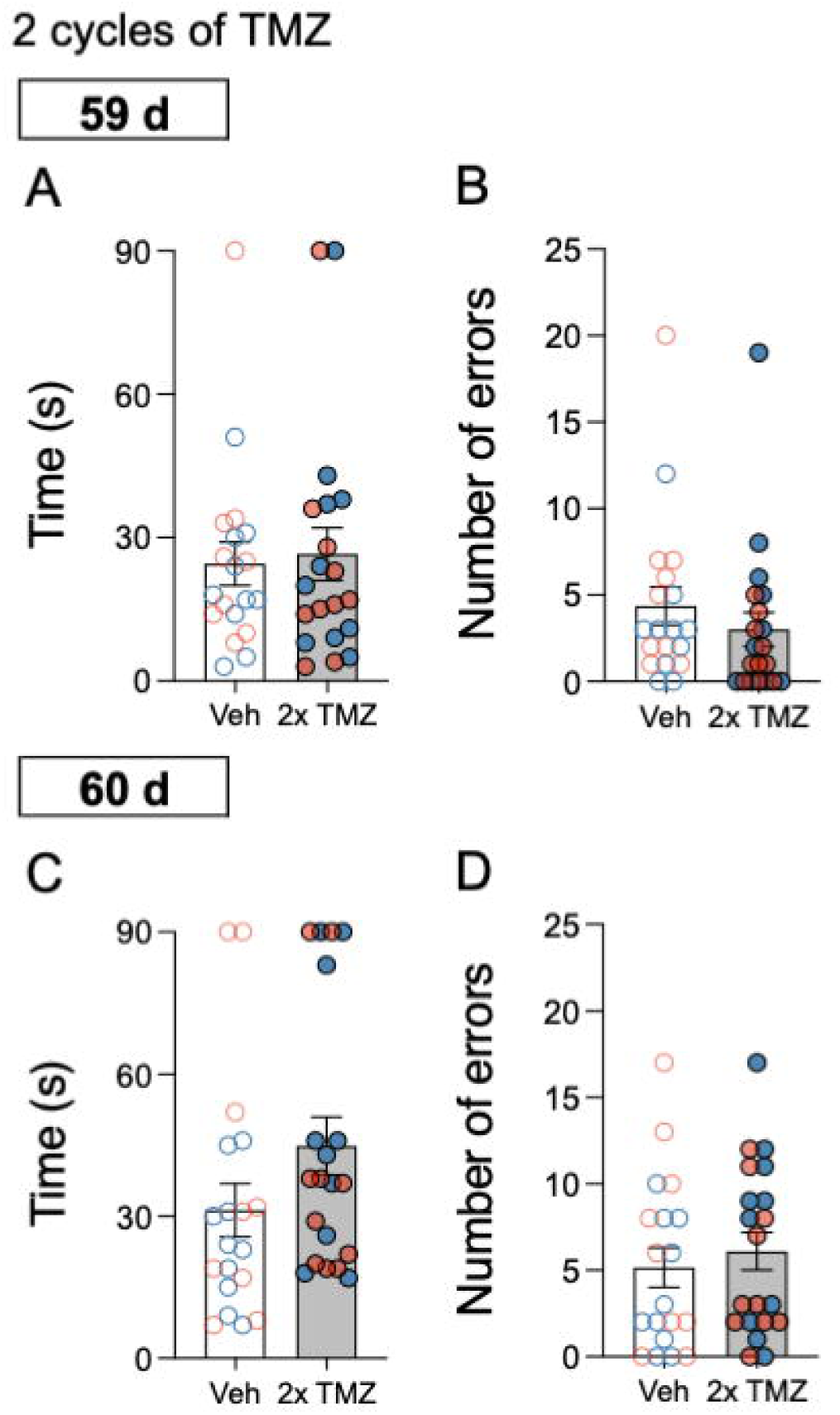
Long-term effects of temozolomide (TMZ) on cognitive performance in a mix-sex cohort of adult rats. **A, C** Changes in time (s) and **B, D** number of errors induced by two TMZ cycles (2x TMZ, 10 days total with 2 resting days in between them, 25 mg/kg, x1/day, i.p.) or vehicle (Veh) as measured 59- or 60-days post-treatment in the Barnes maze test. Data represent mean ± SEM of the each of the defined measurements. Individual values are shown for each rat (blue symbols represent male rats while orange symbols females). Data is shown as a mixed-sex cohort combining both sexes. The effect of TMZ treatment was evaluated through Student *t*-tests.

### Temozolomide induced changes in hippocampal neuroplasticity markers

The neurochemical effects induced by temozolomide were ascertained in the hippocampus at the level of several neuroplasticity markers (i.e., neuronal progenitors, FADD and BDNF) 1-day post-2x cycles of temozolomide in a mixed-sex cohort of adult rats (Fig. 5). There were some overall effects induced by treatment (NeuroD and FADD) that will be described in detail below. Specifically, the regulation of hippocampal NeuroD was used as a positive control of treatment response when measured 1-day post-2x cycles of temozolomide treatment (Fig. 5A-B). As expected, and as previously demonstrated by our group (although only for adult male rats; García-Cabrerizo et al., 2020), 2x cycles of temozolomide treatment (10 days) reduced the number of neural progenitors in the hippocampus of a mixed-sex cohort of adult rats (Fig. 5A-B). The main result that 2x cycles of temozolomide induced a significant reduction in NeuroD +cells/mm^2^ (-117 ± 69 s, *t* = 8.04, *df* = 21, \*\*\**p* < 0.001 vs. vehicle-treated rats; Fig. 5A-B) as measured 1-day post-treatment in adult rats. Similar results were observed for FADD protein, since temozolomide decreased its content in the hippocampus (-19 ± 9 %, *t* = 2.10, *df* = 21, \**p* = 0.048 vs. vehicle-treated rats; Fig. 5C, E). Finally, BDNF was not modulated in the hippocampus by temozolomide treatment (Fig. 5D-E).

**Fig. 5.**
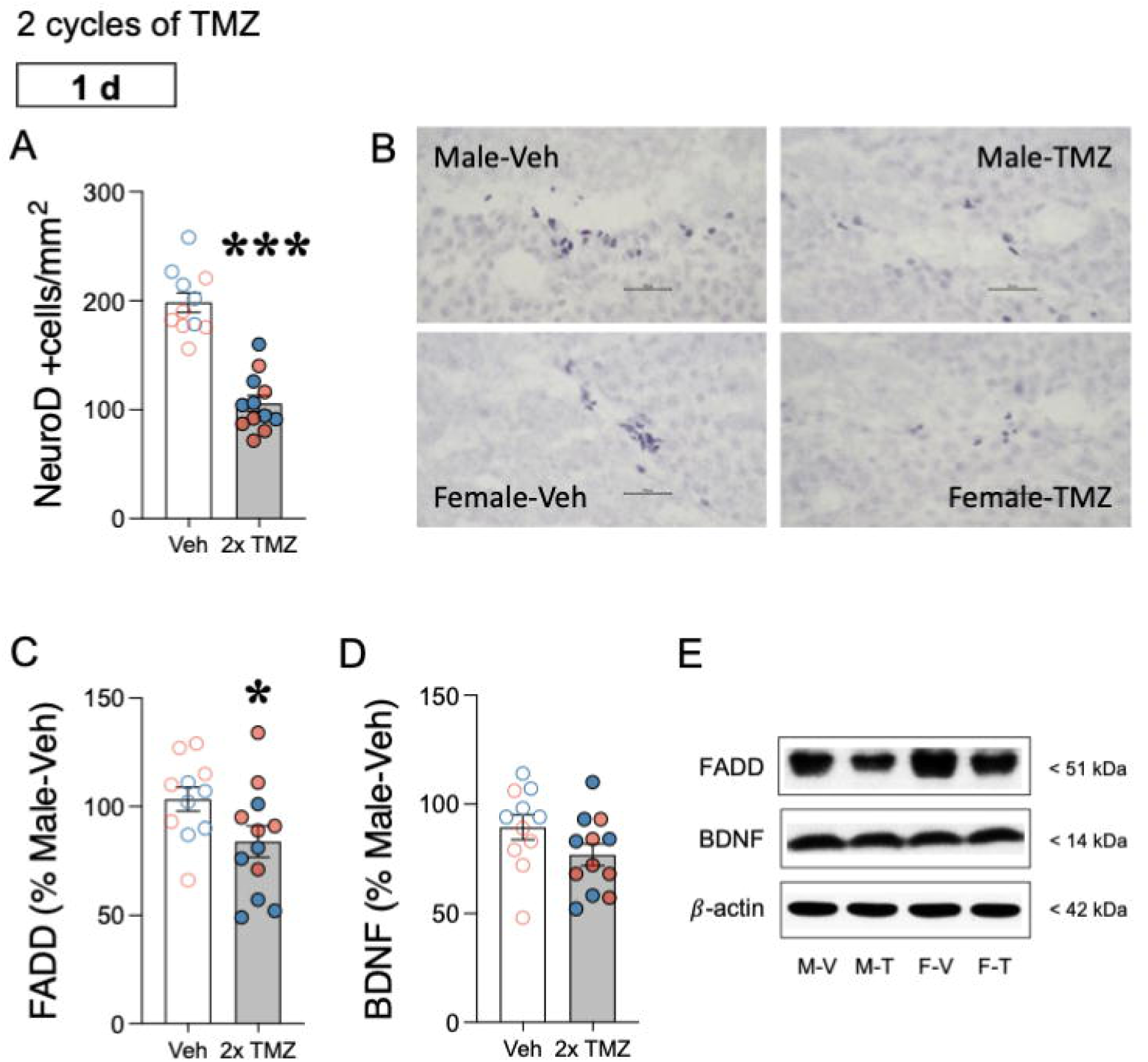
Neurochemical effects of temozolomide (TMZ) in a mix-sex cohort of adult rats. **A** Changes in the number of hippocampal NeuroD +cells/mm^2^ induced by two TMZ cycles (2x TMZ, 10 days total with 2 resting days in between them, 25 mg/kg, x1/day, i.p.) or vehicle (Veh) as measured 1-day post-treatment by immunohistochemistry. Data represent mean ± SEM of the number of NeuroD +cells analyzed as corrected by the area of analysis (mm^2^). Individual values are shown for each rat (blue symbols represent male rats while orange symbols females). Data is shown as a mixed-sex cohort combining both sexes. The effect of TMZ treatment was evaluated through Student *t*-tests (\*\*\**p* < 0.001 when comparing 2x TMZ vs. Veh-treated rats). **B** Representative images of NeuroD +cells (dark blue labeling in a lighter blue granular layer background) for each treatment group (Veh vs. TMZ) for a male or female adult rat as taken with a light microscope using a 63× objective lens. Scale bar: 30 μm. **C** Changes in FADD and **D** BDNF protein levels induced by two TMZ cycles (2x TMZ, 10 days total with 2 resting days in between them, 25 mg/kg, x1/day, i.p.) or Veh as measured 1-day post-treatment by western blot analyses. Data represent mean ± SEM of the protein levels (%Male-Veh). Individual values are shown for each rat (blue symbols represent male rats while orange symbols females). Data is shown as a mixed-sex cohort combining both sexes. The effect of TMZ treatment was evaluated through Student *t*-tests (\**p* < 0.05 when comparing 2x TMZ vs. Veh-treated rats). **E** Representative immunoblots depicting FADD, BDNF, and β-actin for each treatment group (Veh vs. TMZ) for a male or female adult rat (M: male; V: Veh; F: female; T: TMZ).

## Discussion

The main results of this study demonstrated that temozolomide induced antidepressant-like responses in a treatment-duration manner while decreased hippocampal FADD, a neuroplastic marker previously associated with the acute and repeated actions of most antidepressants in a mixed-sex cohort of adult rats. We propose that these unexpected findings break the prior dogma linking increased hippocampal neurogenesis with antidepressant-like efficacy, since it proves that a treatment paradigm that also impaired this neuroplasticity process still induced an antidepressant-like response, in a well-validated screening assay such as the forced-swim test. Therefore, other mechanisms of action, such as the one described through the neuroplastic molecule FADD, might be responsible for the antidepressant-like actions of temozolomide, even in the presence of impaired neurogenesis. Also, we proved no persistent effects on cognitive performance, and all of the experiments were done in a mixed sex-cohort of rats, as opposed to most of the prior literature which mainly centered exclusively in male rodents.

Extensive research in rodents indicated that temozolomide lacked antidepressant-like properties; rather, its administration might be linked to depressive- and anxiety-like behaviors (e.g., Egeland et al., 2017; Pereira-Caixeta et al., 2018; Gan et al., 2019; Dey et al., 2020; Cabrera-Muñoz et al., 2023). This is believed to happen through its primary mechanism of action as a chemotherapeutic agent, which also impairs the proliferation of novel brain cells (i.e., hippocampal neurogenesis; Garthe et al., 2009; Nokia et al., 2012; Egeland et al., 2017; García-Cabrerizo et al., 2020), especially since depression- and anxiety-like behaviors have been related with low levels of this process (Duman et al., 1999; Revest et al., 2009; Kheirbek et al., 2012). In fact, many preclinical studies employed temozolomide as a pharmacological tool to inhibit hippocampal neurogenesis and ascertain its role in various behavioral changes (e.g., Brozka et al., 2017; Cuartero et al., 2019; Luján et al., 2019), including antidepressant-like efficacy (e.g., Gan et al., 2019; García-Cabrerizo et al., 2020; Chong et al., 2021). Our previous results (García-Cabrerizo et al., 2020) suggested in line with other data, that while there is a requirement for adult neurogenesis in mediating some of the beneficial effects of antidepressants (Santarelli et al., 2003), decreasing neurogenesis alone is not sufficient to drive a depressive-like phenotype (Vollmayr et al., 2003; Samuels and Hen, 2011; see further discussion in García-Cabrerizo et al., 2020). More so, the present results showed that temozolomide induced an antidepressant-like response while hippocampal neurogenesis was impaired, an effect that was dependent on the number of treatment cycles received; while 1 cycle (5 consecutive days of treatment) was inefficacious, 2 cycles (10 days administered in 5 consecutive days per week for 2 weeks) demonstrated efficacy. When revising the existing literature, most of the prior studies evaluated different number of temozolomide cycles, normally given as 1 cycle per week, for different durations in length (i.e., from 2 weeks to up to 6 months; see for example Egeland et al., 2017; Gan et al., 2019; Dey et al., 2020; Cabrera-Muñoz et al., 2023), and were done mainly in male mice as opposed to the present study that included a mixed sex-cohort of adult rats. We argue that the discrepancy reported might be related to the accumulative dose received across time, with no effects with just one cycle, efficacy with two, as described in this study, and no further effects or even deleterious ones with prolonged treatment regimens lasting from weeks up to several months (e.g., Egeland et al., 2017; Strokotova et al., 2023). Alternatively, other factors varying across different studies might be contributing to the disparities observed in the results, such as for example sex (just including males, just females or both), age (younger vs. older rodents), species under study (mice vs. rats), the behavioral tests utilized for screening the response (forced-swim test as a predictive test of antidepressant effects vs. other tests more centered in measuring anxiety-like behavior), as well as the time at which screening was done after treatment. Overall, our results in conjunction with the prior data, suggested cycle- and/or length-dependent treatment effects in terms of temozolomide’s antidepressant- vs. depressant-like profile.

The regulation of hippocampal NeuroD was used as a positive control of treatment response when measured 1-day post-2x cycles of temozolomide treatment. As expected, and as previously demonstrated by our group (although just in adult male rats; García-Cabrerizo et al., 2020), 2x cycles of temozolomide treatment (10 days) reduced the number of neural progenitors in the hippocampus of a mixed-sex cohort of adult rats. Interestingly, this is the same time point of analysis at which temozolomide induced antidepressant-like effects in the forced-swim test, suggesting a dysregulation between these two processes. At this point, the results pointed out at the probable regulation of other neuroplastic mechanisms rather than neurogenesis or BDNF (which was not altered by temozolomide in line with prior studies; Pereira-Caixeta et al., 2017), to justify the observed antidepressant-like efficacy induced by temozolomide. In this regard, we explored hippocampal FADD protein as a potential neuroplasticity target that could mediate temozolomide’s antidepressant-like response, since we previously demonstrated that FADD was down-regulated by many antidepressants with different mechanisms of action, and that lower levels of FADD protein correlated with decreased immobility in the forced-swim test as induced by desipramine (García-Fuster and García-Sevilla, 2016). Our results showed indeed that temozolomide (2 cycles) decreased hippocampal FADD protein in a mixed sex-cohort of adult rats as measured 1-day after treatment at the same time the antidepressant-like response was scored. Unfortunately, no correlation analyses could be run with the behavioral results since rats exposed to the behavioral characterization were from a different study than the ones where neurochemistry was ascertained. Since decreased FADD could participate in the neuroplastic actions activated by many antidepressants (García-Fuster and García-Sevilla, 2016), we propose that it also does so following this short treatment regimen of temozolomide. Aligned with our data, temozolomide also showed some beneficial effects on neurocognitive functioning and affective-like responses (i.e., improved anxiety in the open field and elevated plus maze) when administered in rats with induced glioblastoma, which were also independent of neurogenesis and regulated via other routes, such as oxidative stress or reduced cytokines for instance (see more details in Moslemizadeh et al., 2022). Altogether, this data together with the prior literature suggested that other molecular mechanisms might be mediating the response induced by temozolomide, ensuring the need for further studies.

The last part of the study aimed to ascertain whether our experimental paradigm of temozolomide would affect long-term cognitive performance in the Barnes maze, since prior literature seemed contradictory in this regard. On the one hand, some studies suggested that the depletion of new neurons might be a key mechanism associated not only with the development of depression-like symptoms but that might also affect hippocampus-dependent learning memory, and novelty processing (e.g., Pereira-Caixeta et al., 2018). However, other studies suggested quite the opposite; reducing hippocampal neurogenesis with temozolomide improved spatial reversal learning in rats (Brozka et al., 2017) or improved cognitive impairment after stroke in mice (Cuartero et al., 2019). To add more inconsistency and contrary to any of the detrimental or beneficial effects reported, we now showed that 2 cycles of temozolomide preserved the cognitive function of a mixed sex-cohort of adult rats up to 60 days post-treatment. These discrepancies aligned with a report that suggested that the effects of hippocampal neurogenesis suppression on spatial learning might be age-dependent; while suppression of neurogenesis during juvenility altered hippocampal development and impaired learning, pre-existing neurons might have compensated for the lack of new hippocampal neurons during adulthood and aging, since no effects were observed on spatial learning (Martinez-Canabal et al., 2013).

## Conclusion

Against previous data suggesting that the depletion of new hippocampal neurons is a key mechanism associated with the development of depression-like symptoms, the present study demonstrated that temozolomide decreased hippocampal neuronal progenitors while still induced antidepressant-like effects in a mixed-sex cohort of adult rats. This effect was dependent on treatment duration and on the test used to score the response and was probably mediated through FADD protein, a neuroplastic marker previously associated with the acute and repeated actions of several antidepressants. Therefore, we argue that any mood changes associated with temozolomide treatment for glioblastoma might be associated with treatment duration together with the concomitant manifestation of depressive-like behaviors. In fact, the data suggested that short-duration treatments of temozolomide could be beneficial for such comorbidities. Finally, we propose FADD as a key neuroplastic marker associated with the potential beneficial effects of temozolomide in a mixed sex-cohort of adult rats.

## Data availability statement

The datasets used and/or analyzed during the current study can be made available from the corresponding author on reasonable request.

## Ethics statement

The animal study was approved by the studies involving animals were reviewed and approved by the Local Bioethical Committee “Comité de Ética de Experimentación Animal” and by the Regional Government (approved protocol number SSBA 21/2024 AEXP). The study was conducted in accordance with the local legislation and institutional requirements.

## Author contributions

**LG-M:** Data curation, Formal Analysis, Methodology, Investigation, Writing-review and editing. **MG-F:** Conceptualization, Data curation, Formal Analysis, Supervision, Resources, Original Draft Preparation, Writing - Review & Editing.

## Acknowledgments

The authors would like to acknowledge the procedural assistance provided by Neus Mateu Mercader (TECH-2023 from «CONSCIENCIA IdISBa: consolidar la ciència IdISBa», funded by ITS2023-057).

## Supplementary Materials

Supplementary material associated with this article can be found, in the online version.

