## Supplemental Materials for "Unexpected antidepressant-like effects of temozolomide in a mixed sex-cohort of adult rats: role of hippocampal FADD protein"

### Supplementary Materials

**Fig. S1. Basal performance in the forced-swim test prior to any drug treatment.** Data represents the time spent (s) immobile, climbing or swimming in the forced-swim test as performed prior to any drug treatment for male and female rats. These basal values were used to counterbalance rats in different treatment groups as detailed in Fig. 1A.

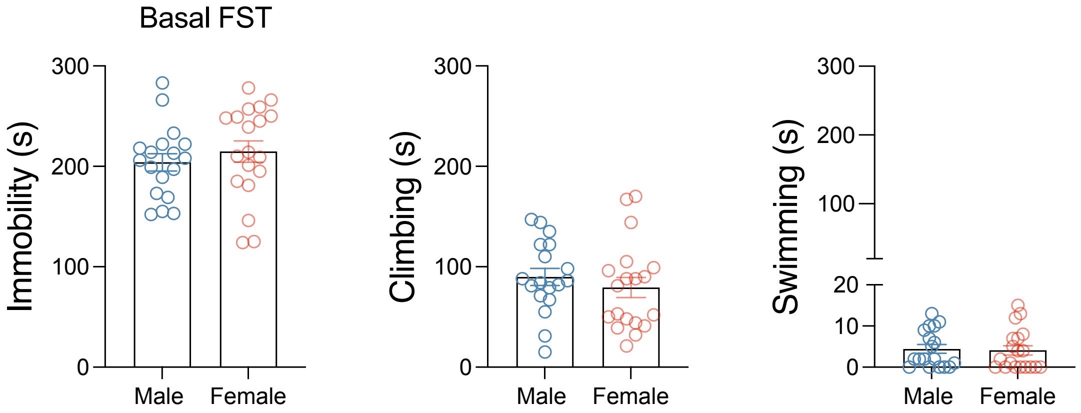
